# Multiparameter cell characterization using nanofluidic technology facilitates real-time phenotypic and genotypic elucidation of intratumor heterogeneity

**DOI:** 10.1101/457010

**Authors:** Kristin G. Beaumont, Wissam Hamou, Nenad Bozinovic, Thomas R. Silvers, Hardik Shah, Arpit Dave, Kimaada Allette, Maya Strahl, Ying-chih Wang, Hanane Arib, Alesia Antoine, Ethan Ellis, Melissa Smith, Brandon Bruhn, Peter Dottino, John A. Martignetti, Eric Schadt, Mark White, Robert Sebra

## Abstract

Genetic and functional complexity of bulk tumor has become evident through rapid advances in sequencing technologies. As a unique integrated approach to characterizing tumor heterogeneity, we demonstrate the multifaceted capabilities of a novel nanofluidic platform to enable single-cell phenotypic and genetic profiling of ovarian cancer patient-derived tumor cells. This approach has enabled increased resolution of tumor cell phenotypic and genetic heterogeneity, providing a better understanding of underlying biological drivers of the disease. A range of CA-125 expression levels is observed within cells from individuals, demonstrating clonal diversity consistent with other phenotypic data. Further, TP53 mutation analysis demonstrates a sub-population of cells exhibiting high mutation frequency that likely drives downstream growth kinetics and protein expression. Finally, genomic data is orthogonally used to address clonal heterogeneity across ovarian tumors when compared to bulk sequencing, illustrating the potential for single-cell sequencing data integrated with cellular functional and growth data toward future therapeutic intervention.

## INTRODUCTION

Single cell sequencing analysis is a rapidly emerging field that has only recently been applied to identifying the diverse cellular landscape that constitutes living tissue and new, innovative technological advances have enabled progress in this field.^1^^-^^3^ This approach has great potential to reveal previously obscured complexity in a variety of heterogeneous biological systems, but presents technological challenges that are unique to the single cell vantage point. One hurdle that needs to be overcome in single cell sequencing is capturing single cells that are 1) of high enough quality to yield sequenceable material and 2) numerous enough to yield population-representative data.

Historically, laser dissection and FACS have been used to isolate single cells for sequencing.^4^ Laser dissection provides user-directed specificity in target cell selection, but is highly limited in the number of cells (tens to hundreds) that can be isolated. FACS also offers marker-based specificity during selection and permits the isolation of hundreds or thousands of cells, but can be time consuming (often resulting in sample degradation) and requires labor-intensive downstream processing for sequencing. Droplet-based single cell sequencing methods are the most recent innovative approaches that facilitate the isolation and sequencing of large numbers (thousands) of single cells in a highly efficient manner. These methods combine single cell isolation (via Poisson loading of single cells into fluid droplets) with cell lysis. This approach is somewhat limited in scope as it does not permit real time selection of target cells and is destructive to cellular material.^5^^,^^6^ Moreover, Poisson encapsulation is inherently inefficient and throughput-reducing, resulting in mostly empty droplets. Throughput of single cell approaches has been greatly improved by the addition of cell barcodes for multiplexing^7^ and microfluidics to improve capacity and handling efficiency. Platforms such as Chromium, ddSeq and C1 now combine microfluidic devices and cell barcoding to enable cDNA generation from hundreds to thousands of single cells at once.

Finally, as single cell sequencing technologies develop, investigators can ask increasingly challenging research questions that require integration of complex cellular phenotype data with sequencing data. Approaches such as CITE-seq, where markers for cellular protein expression are combined with Drop-seq based cell isolation and processing, have begun to address these challenges, allowing sequencing information to be linked to characterized cells.^8^ However, the types of phenotypic data accessible by this method are currently limited to antibody-labeled surface protein expression and cannot include complex metrics such as growth kinetic or cellular function.

To our knowledge, this study demonstrates the first platform to support characterization linking phenotype to genotype at the single cell level across a culture system, which will enable a more thorough, integrated analysis of tumor heterogeneity, and could rapidly accelerate the development of precision oncology for many tumor types. For this study, we focus on high-grade serous ovarian cancer (HGSOC) as a model to demonstrate the key components of our approach in providing a potentially more sensitive means of characterizing tumor heterogeneity at single cell resolution.

The poor survival rate of ovarian cancer, the most lethal female gynecologic malignancy which will be diagnosed in almost 200,000 women worldwide this year,^9^ remains relatively unchanged over the past four decades.^10^ HGSOC will account for nearly 80% of deaths from this disease. Though complete clinical response to surgical debulking and platinum-based chemotherapy is initially observed, three out of four women will experience tumor recurrence and succumb to platinum-resistant disease within five years of initial presentation.^11^ Recent progress in precision oncology suggests the ability to move beyond the currently stagnant universal treatment assumptions for HGSOC and towards a more personalized approach based on tumor genotyping to reveal optimal therapies. Treatment options can include traditional cytotoxic chemotherapeutics and targeted agents aimed at providing the greatest antitumor response while also improving survival and, ideally, maintaining a higher quality of life. One major limitation to delivering precision oncology in HGSOC, and other cancers, is the current inability to account for the degree of temporal and spatial heterogeneity present within tumors.^12^ For example, patients with chronic lymphocytic leukemia who have detectable subclonal driver mutations prior to initial treatment had earlier recurrences and shorter survival times.^13^ Similarly, quantitative assessments of intratumor heterogeneity in HGSOC may be predictive of survival following chemotherapy.^14^

Drug treatment itself has also been shown to drive sub-clones into distinct tumor populations, as shown with acute myeloid leukemia^15^ and multiple myeloma.^16^ Continuing to analyze HGSOC tumors *en masse* or even from multiple slices is unlikely to identify adequately and describe driver mutations and actionable pathways with direct consequences on a tumor’s response to treatment.^12^^,^^17^^-^^19^ For example, The Cancer Genome Atlas (TCGA) analysis of HGSOC has identified TP53 mutations as the predominant molecular feature of this disease, albeit with a relatively “long tail” of rare but potentially actionable mutations.^20^ Other analyses of HGSOC tumors have also documented a high degree of intratumoral and spatial heterogeneity.^14^^,^^21^^-^^24^ Unfortunately, currently available HGSOC functional assays from patient-derived materials may not appropriately model this clonal complexity.^25^

This work presents proof-of-concept data that demonstrates the capabilities of a new nanofluidic cell analysis system to facilitate single cell manipulation, culture, assay and analysis, and test its application across multiple modalities. We have chosen several representative cell types (including those derived from immunological and cancer lineages) in addition to cells derived from HGSOC patient tumor biopsies to illustrate the flexibility of this approach. Relying on light-induced dielectrophoresis technology, we interrogate tumor heterogeneity using single tumor cell sorting with subsequent analysis of growth kinetics and protein expression, followed by single cell genetic profiling of HGSOC patient-derived tumor cell lines.

## RESULTS

### Single Cell Handling Technology Overview

All data were collected using a novel, flexible nanofluidic platform, (Berkeley Lights, Inc. (BLI)), for single cell and batched multicellular selection and manipulation using light-induced dielectrophoresis. This system (Beacon^TM^ prototype platform) is overviewed in Figure 1A. The platform integrates mechanical, fluidic, electrical and optical modules, enabling single cell manipulation, culture and imaging with a micrometer level of precision. The primary strength of the system is the disposable nanofluidic device with 1350-3500 individual chambers (pens) depending on chip design, each capable of holding sub-nanoliter volume. The fluidics subsystem delivers media to the chip for cell culture or reagent delivery and an actuated needle automates import and export of cells from incubated well plates. A 3-axis robotic stage enables imaging of the entire chip area at 4x or 10x magnification in both brightfield and fluorescent modes for orthogonal characterization of morphologic, functional, and biochemical events. White light combined with a digital micromirror device (DMD) array is used to structure desired patterns for light-actuated dielectrophoresis or fluorescent illumination.

**Figure 1.**
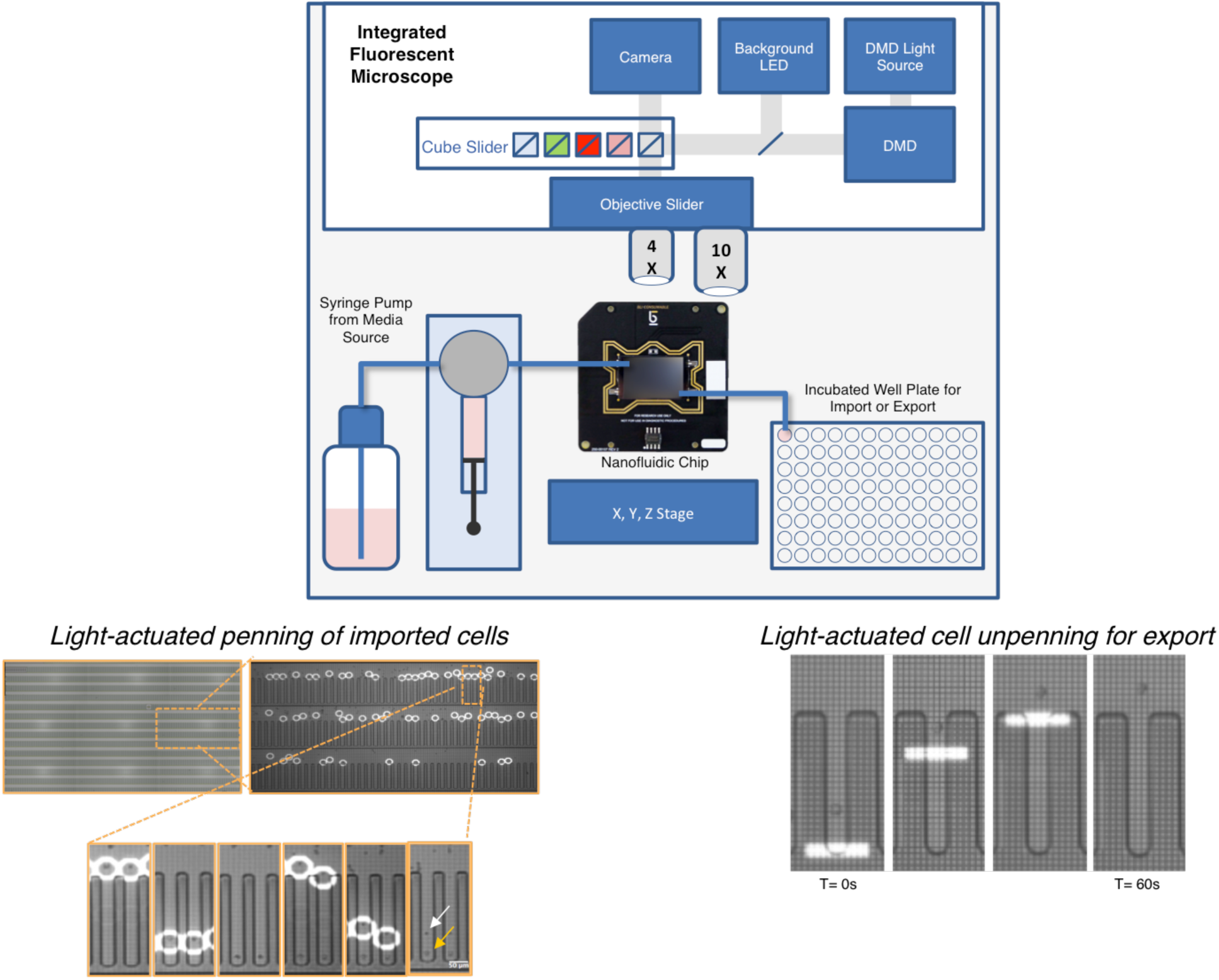
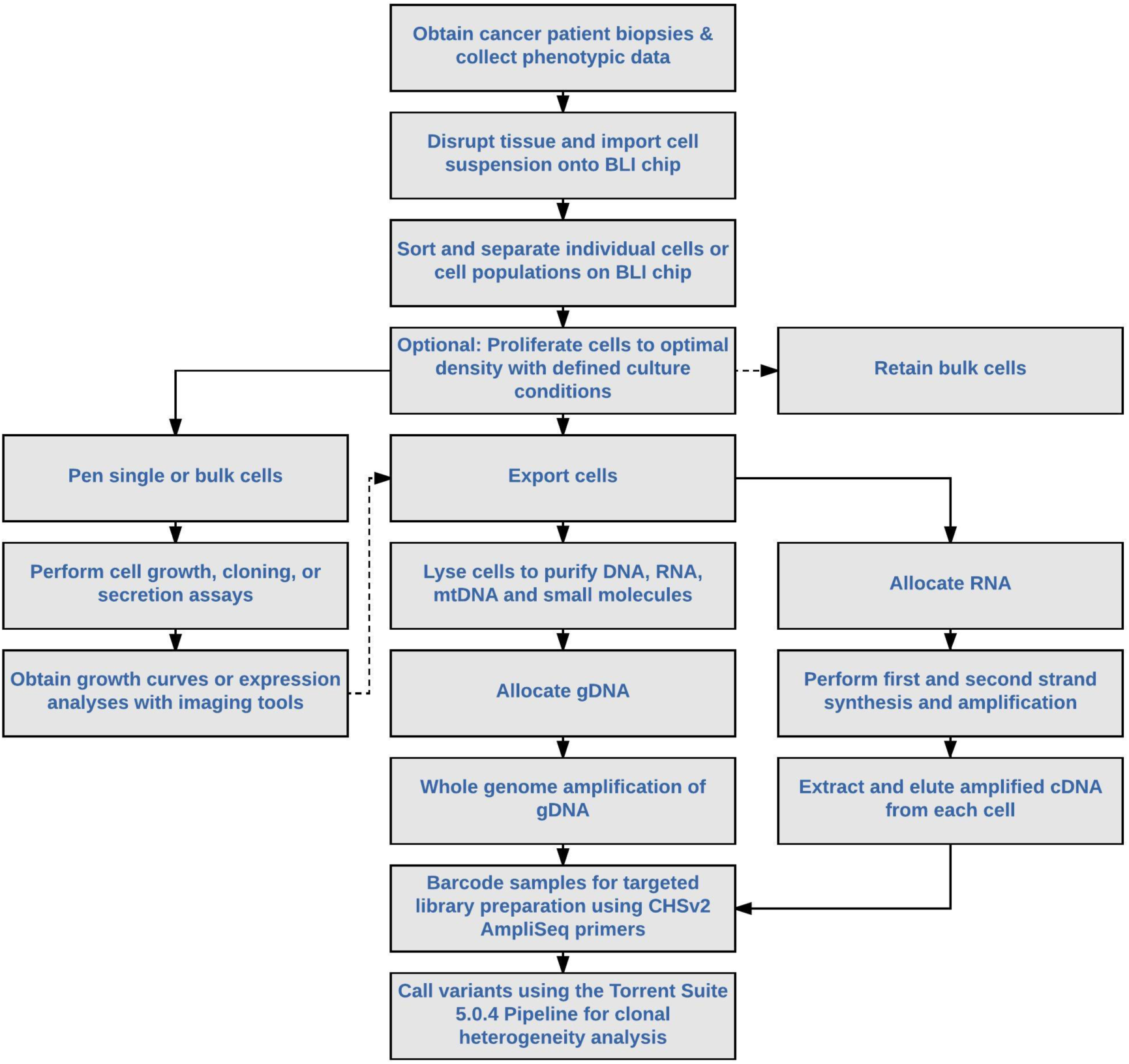
Overview of automated nanofluidic cell sorting, analysis platform and workflows. **(A)** Cell samples in a 96 well plate are placed in an environmentally-controlled well plate incubator within the BLI Beacon^TM^ prototype platform. A needle on an automated Z stage dips into the well containing cells and a syringe pump connected to the OptoSelect^TM^ chip and tubing aspirates a sample from the plate. An integrated fluorescent microscope creates light patterns using a digital micromirror device (DMD) array while a background light-emitting diode (LED) light source illuminates the field of view for a camera to image live processes. Assay reagents are loaded via the 96 well plate with accompanying well plate incubator and a larger media bottle is used to facilitate perfusion-based cell culture over multiple days. **(Left inset)** Composite image of the central 9 fields of view on the nanofluidic chip showing representation of annotation and isolation of single cells using BLI’s OEP technology. Using machine vision and automated path planning, single OVCAR3 cells are selected and isolated into individual 50-µm wide pens. After flushing unpenned OVCAR3 cells from the chip, Jurkat cells are imported into the chip and a single Jurkat cell (white arrow) is placed into each pen containing a single OVCAR3 cell (orange arrow). **(Right inset)** A single Jurkat cell is unpenned using OEP prior to export into a 96-well plate for storage or additional molecular and functional analysis. Images of single cells after unpenning are taken before the syringe pump pushes media through the chip to flush the cell into the well plate **(B)** Overview of the workflows employed for analysis of primary patient-derived tumor tissue.

The silicon base of the nanofluidic chip is composed of approximately 1 million light-actuated switches that activatea dielectrophoretic force and repel particles resulting in light “cages” that trap and move desired particles, including cells and beads, at 5-20 μm/s. This light-induced force enables OptoElectroPositioning (OEP) where cells and beads can be positioned into the sub-nanoliter volume chambers on demand. Patterned chip surface chemistry and overall hydrophilicity can be modified such that the surface either promotes or prevents adhesion. The experimental platform is unique amongst other single cell technologies as it only uses OEP force for cell selection and employs fluidics for cell transport resulting in little-to-no damage to cell viability, which is critical to enable multiple downstream applications, including genomic analysis as well as on- and off-chip functional characterization of selected cells.

### Cell import, penning and export

Using this platform, we are able to load, position, culture and export cells in a defined and trackable manner. Target cells are specifically placed into numbered pens, where their growth is characterized and the cells are phenotyped, prior to export from the platform into a tracked position of a well plate for further processing and sequencing. As an overview, cells are imported from an incubated well plate onto the nanofluidic chip and guided into pens using light-induced OEP force. Cells can be placed into pens (as single or defined groups of cells) in a highly parallel fashion. One key advantage of the nanofluidic chip design that enables simultaneous and independent interrogation of many different cells is the sequestration of pens from each other. Following cell import, the main channel is flushed to eliminate any unwanted cells in the fluidic path; cells in the pens are fluidically isolated from the channel but remain accessible to soluble factors from the bulk media by diffusion.

An example of a complex penning strategy is shown in Fig. 1A, in which a single OVCAR3 cell is positioned in a pen, followed by a single Jurkat cell (Fig. 1A left inset). In this case, 31 images were acquired to cover the entire chip area and a sampling of 9 stitched images centrally located on this chip design is shown. A machine vision algorithm is used to identify cells based on their size and morphology, and desired cells were shepherded by OEP into target pens on the chip (Fig. 1A left inset - time lapse). Cells can be distinguished based on circularity, diameter or fluorescent intensity (in DAPI, FITC, Texas Red or Cy5). Following culture and characterization, these same cells can be exported off of the nanofluidic chip for sequencing or further processing. The export process is the reverse of the cell placement process (shown for a Jurkat cell in Fig 1A right inset), where cells from individual or groups of pens are exported using OEP and then gently flushed into individual, tracked wells of an external 96- or 384-well plate in separate 2 µL volumes. A frame-by-frame history of cells, unpenned and ready for export, is automatically archived for additional analysis.

Fig. 1B illustrates the utility of this platform in context of a complete tumor heterogeneity study, intended to link cellular phenotype with genotype. The workflow begins with patient biopsy collection, disaggregation of tissue, followed by cell sorting, monitored *in situ* cell growth, protein expression analysis and genomic analysis.

### Rare cell detection and automated cell annotation

This approach can be easily adapted to facilitate universal or discriminate selection of cells based on morphology and fluorescence across a broad range of inputs, minimizing dependence on sample input criteria and can be especially useful where cells of the desired phenotype are rare relative to the total number of cells in a complex sample. Specifically, we have developed penning strategies that permit inputting cell suspensions as dense as 2×10^7^ cells/mL, which translates to 50,000 cells loaded into the chip at once. By performing sequential importing, penning, and flushing cycles, we can robustly sample hundreds of thousands of cells. Further improvements to the instrument have enabled automation that increases this sampling capacity. The ability to locate and track cells precisely is critical for dynamic experimentation examining both cell growth and functional biology, and iterative processing could further enable applications such as identifying small numbers of circulating tumor cells in patient blood samples. As a case study, we demonstrated the ability to select as few as 1 bead in 100,000 cells for penning by fluorescence (Fig. 2). We spiked known numbers of green fluorescent microspheres into a background of unstained Jurkat cells, loaded onto a nanofluidic chip and imaged by both brightfield and fluorescence (Fig. 2A). Using segmentation and intensity threshold image processing algorithms embedded in the system software, we were able to identify the number and X,Y-coordinates of cells and beads in each field of view (Fig. 2B) and used this information to calculate bead concentration. Correlation analysis revealed a χ^2^=0.99 between the measured spiking ratio and the observed bead concentration (Fig. 2C).

**Figure 2.**
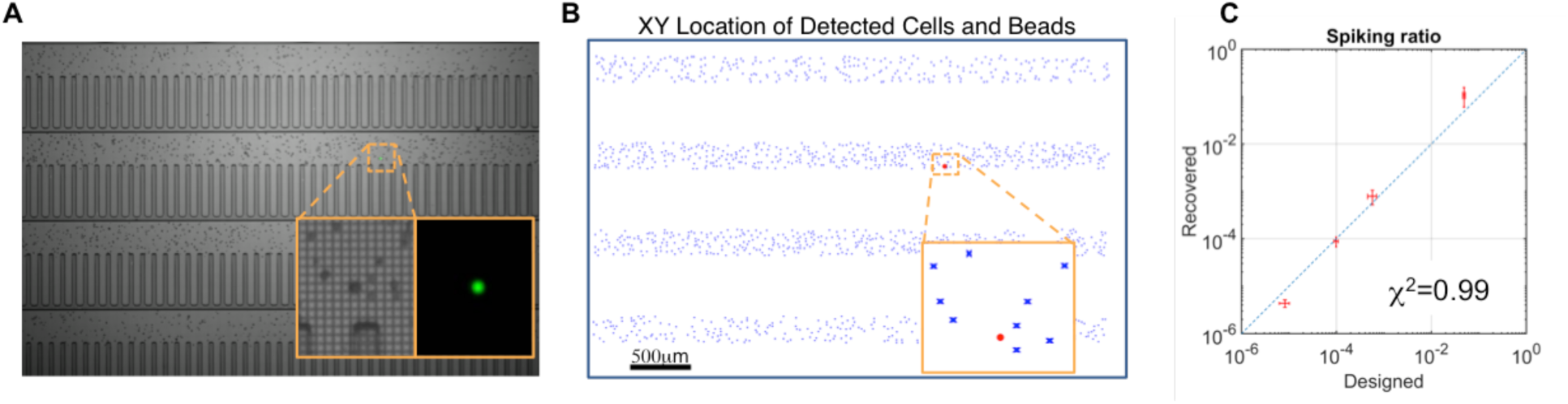
Rare cell detection demonstration. **(A)** Identification of rare cells is performed with fluorescent imaging. Fluorescent green beads are spiked into a suspension of Jurkat cells at various frequencies and are identified by fluorescent imaging (inset). **(B)** XY coordinates of each cell captured by the machine vision algorithm and the rare fluorescent bead (red) can be identified in the non-fluorescent cell background (blue) **(C)** Using the known nanofluidic chip volume and number of cells or beads identified by imaging, we are able to plot the designed frequency vs the recovered frequency in each experiment. The range of designed frequency was a bead to cells ratio of 1:100 (10^-1^) down to 1:100,000 (10^-5^).

In order to track functional characteristics of individual cell populations over time, the experimental platform has integrated four fluorescent imaging channels and automated cell annotation pipelines to monitor stained cells throughout the course of an experiment. Fluorescent signals can be used to inform penning of cells or as a means to track phenotype. This capability is demonstrated in Fig 3A, in which we co-penned two different cell types, A2780 human ovarian cancer cells and peripheral blood mononuclear cells (PBMCs). We stained both cell types with DAPI, MitoTracker Green, and Anti-CD8, PE-CF594. Use of Anti-CD8 allowed discrimination of PBMC subpopulations, such as T cells, based on CD8 surface expression. Penned cells were imaged across the entire chip to extract size and fluorescent intensity measurements using the system software and demonstrate the existence of several distinct clusters of cells based on marker expression (Figs. 3B, C, D, E).

**Figure 3.**
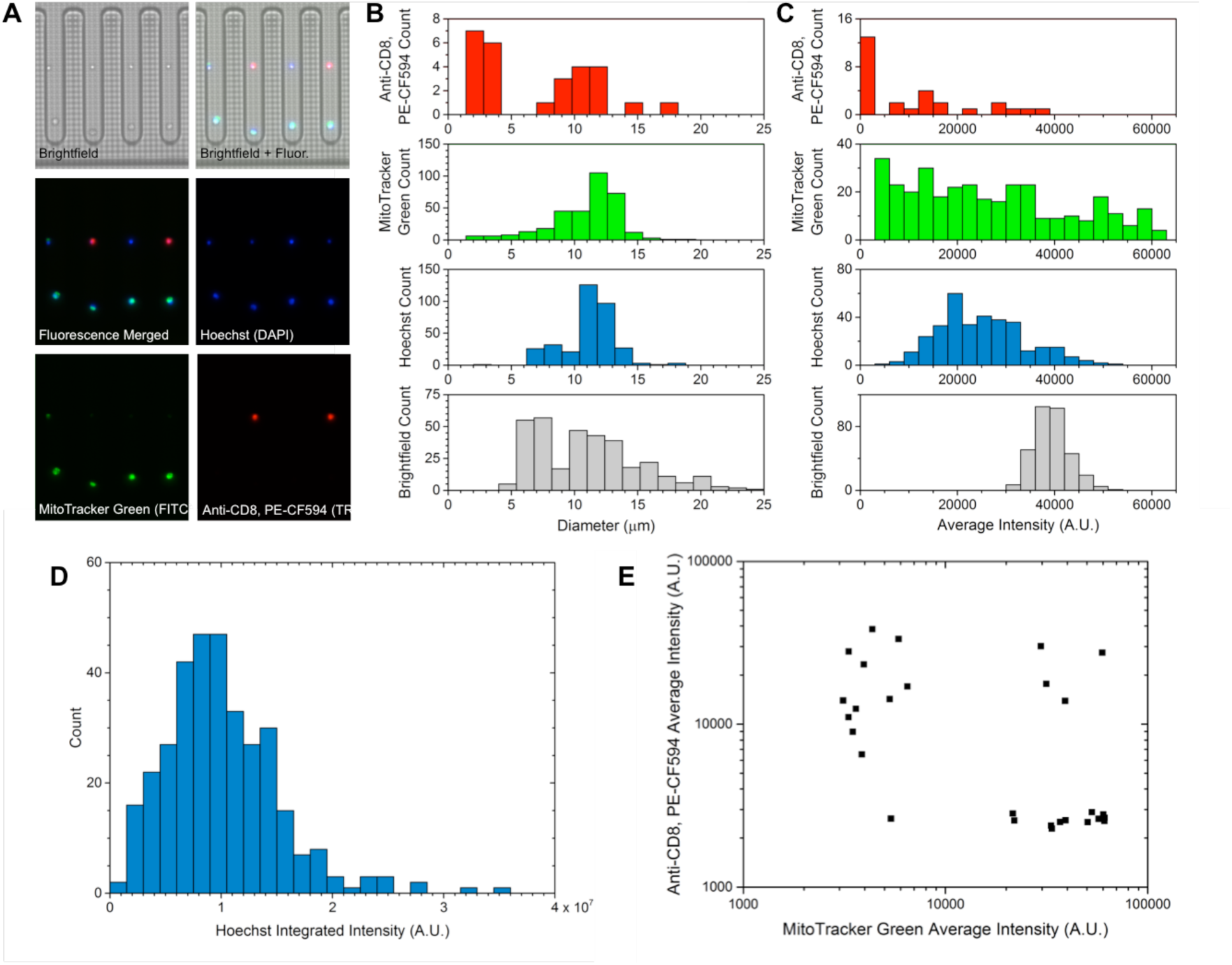
Automated annotation of co-localized cancer and PBMCs. **(A)** Brightfield and fluorescent images of 4 representative pens that each contain a single cancer cell and PBMC as positioned via OEP. Each image is labeled with the channel(s) used for visualization as well as the fluorophore depicted (if applicable). The exposure time was 50 ms for brightfield, 60 ms for DAPI, 400 ms for FITC, and 750 ms for Texas Red. **(B-C)** Histograms showing the area-based diameters and average intensities of all cells on the chip as measured under brightfield and each fluorescent channel. These parameters were obtained via segmentation- and threshold-based image processing algorithms embedded in the system software. **(D)** A histogram showing the integrated intensity of all cells on the chip as measured under DAPI (corresponding to the Hoechst stain). **(E)** A scatter plot depicting the average intensity of all cells on the chip as measured under FITC and Texas Red (corresponding to the MitoTracker Green and Anti-CD8, PE-CF594 stains, respectively). Several distinct clusters or populations of cells are identified.

### Characterization of cell growth

In addition to penning target cells and characterizing cellular phenotypes, cells can be cultured on-chip over the course of several days, which enables the isolation and expansion of cells of interest from polyclonal populations. Here, murine OKT3 cells were penned using OEP and cultured for 4 days. Images were acquired every 120 min for 48 h and machine vision algorithms were used to generate cell counts at each time point that were plotted to characterize growth patterns in MatLab (MathWorks, Inc.).

The mean observed doubling time was 16 h over 48 h of growth with no detectable loss of cell viability – this is comparable to the observed 11-14 h doubling time of this cell line off-chip under standard culturing conditions. A range of growth rates are observed when individual single-cell derived clonal growth curves are plotted over time from the OKT3 cell line (Fig. 4A). Individual pens and cell growth characteristics are shown in Figures 4B and 4C. These data strongly suggested that manipulation by OEP does not impact the viability of cells, and that nutrient transport from the channels to the pens can support expected levels of cell growth over time.

**Figure 4.**
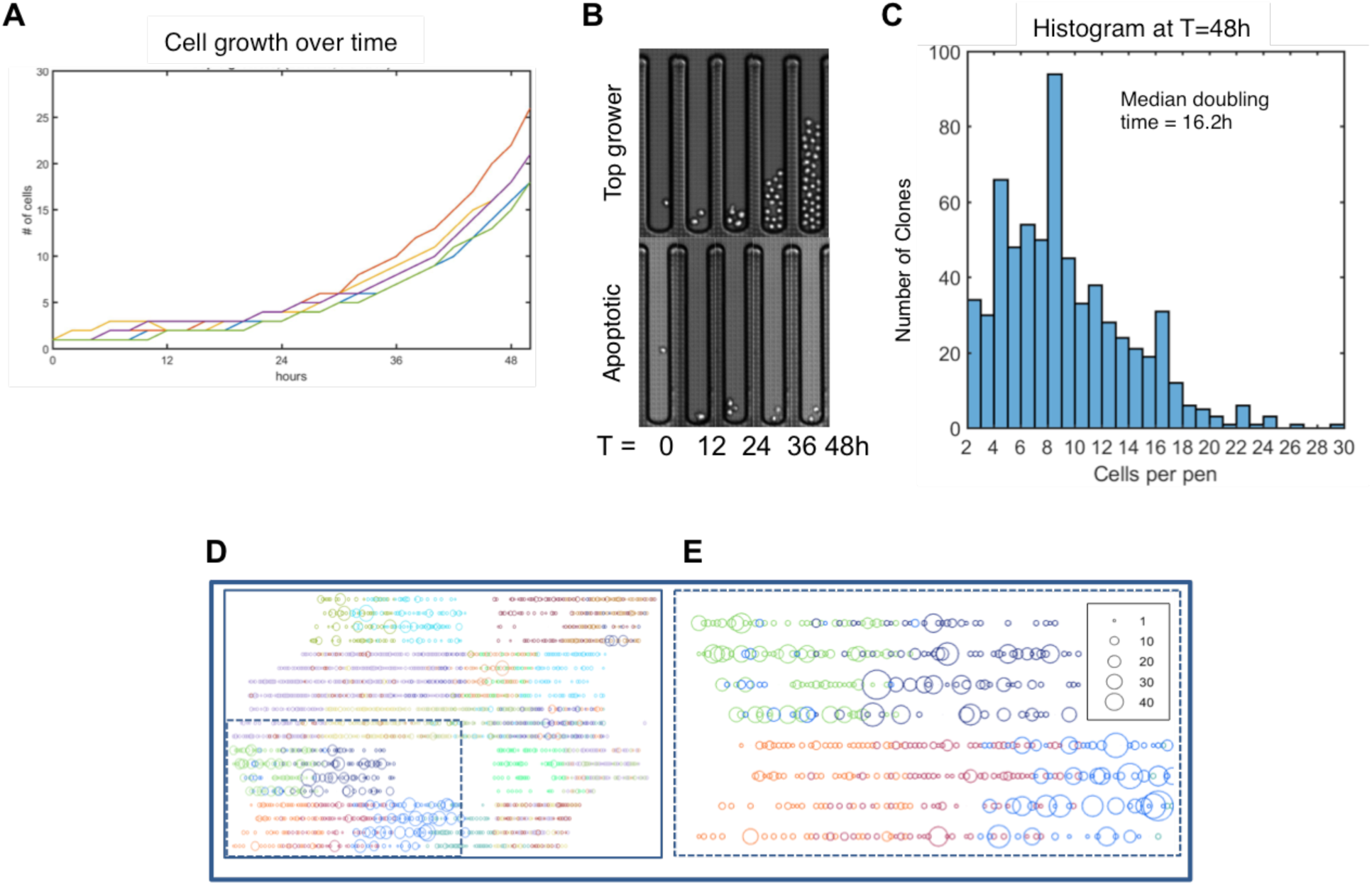
Growth of OKT3 suspension cells cultured on chip. **(A)** Growth curves for 5 selected pens over 48 h **(B)** Time lapse of clone with fastest growth rate (top) and example of identified apoptotic pen (bottom). **(C)** Histogram of cell count at t=48 h. Total number of pens that had two or more cells is 620; 370 single cells failed to divide. Mean growth after 48 h: 8.8 cells. **(D)** Map of multiple OKT3 clones which are independently loaded on the nanofluidic chip and tracked – each sample is represented by a different colored circle and the size of the circle represents the number of cells present for each clone after 48 h. **(E)** Map of inset from panel **(D)**.

The cell penning and tracking strategies described here can also be used to enable investigations with multiple cell populations - many different cell types (or clones) can be penned, grown and characterized on the same chip. Fig. 4D and E show the penning and functional tracking of multiple cellular clones on the same chip, where 4D shows an overview of clone location for the entire chip and 4E shows an example region of interest from this chip, where there is a large growth difference among clones. These data demonstrate both the ability to identify the diversity of cells on the chip, as well as monitor their growth heuristics in real-time, which are automatically tracked by the platform’s integrated computer system.

### Phenotypic analysis of ovarian cancer cells

A REMARK diagram summarizing the patient-derived cell lines used in this study is shown in Fig. 5.

**Figure 5.**
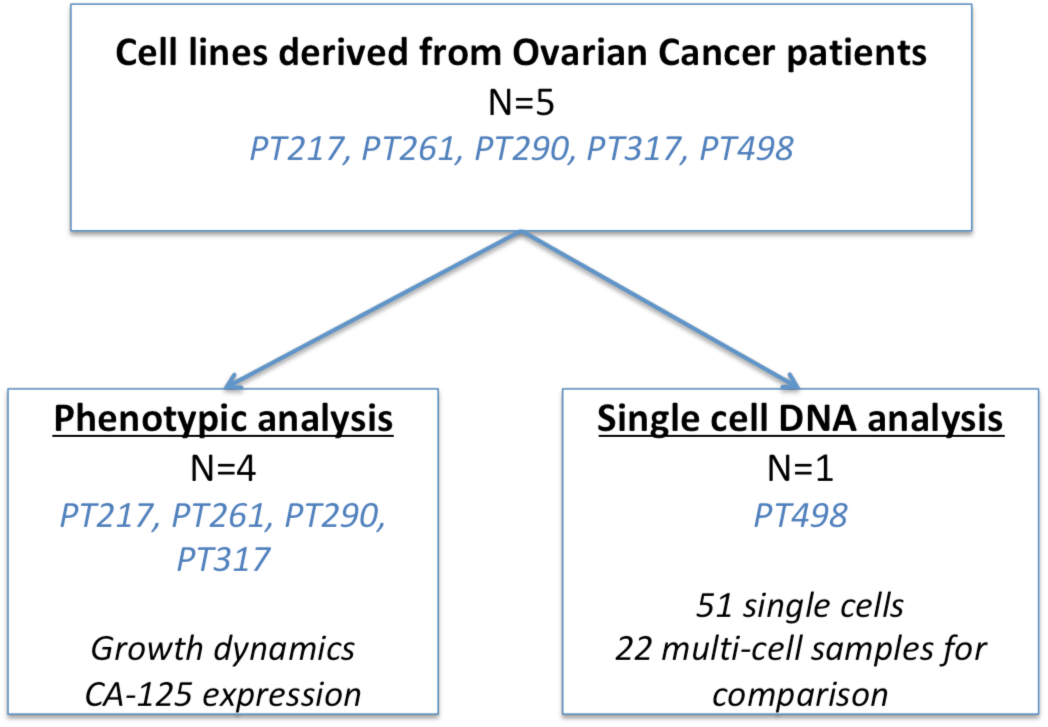
REMARK diagram summarizing patient derived cell lines used in growth, protein expression, DNA and RNA sequencing experiments.

### Growth characteristics

In order to facilitate phenotypic comparison, multiple independent cell populations can be cultured on a single nanofluidic chip, as shown in Fig. 6, where four ovarian cancer patient-derived tumor cell lines (PT217, PT290, PT317, and PT261) and immortalized OVCAR3 cells were grown in parallel (Fig. 6A shows the penning map for these cells, in addition to a brightfield image of cells after loading in one sample region). Cells were penned using OEP, cultured on-chip for six days with image acquisition every two hours, and counted using the platform’s automated cell tracking and counting features (Fig. 6B). From experimental iterations (n=9), we determined that a minimum of 6-10 cancer cells generally need to be penned together on day 1 for survival and growth, possibly due to the presence of cell-secreted factors in the local pen microenvironment. Interestingly, it appears that the cells begin to aggregate at days 2 and 3, eventually forming a multicellular spheroid that could be used for downstream diagnostic and/or prognostic experimentation (Fig. 6C and inset).

**Figure 6.**
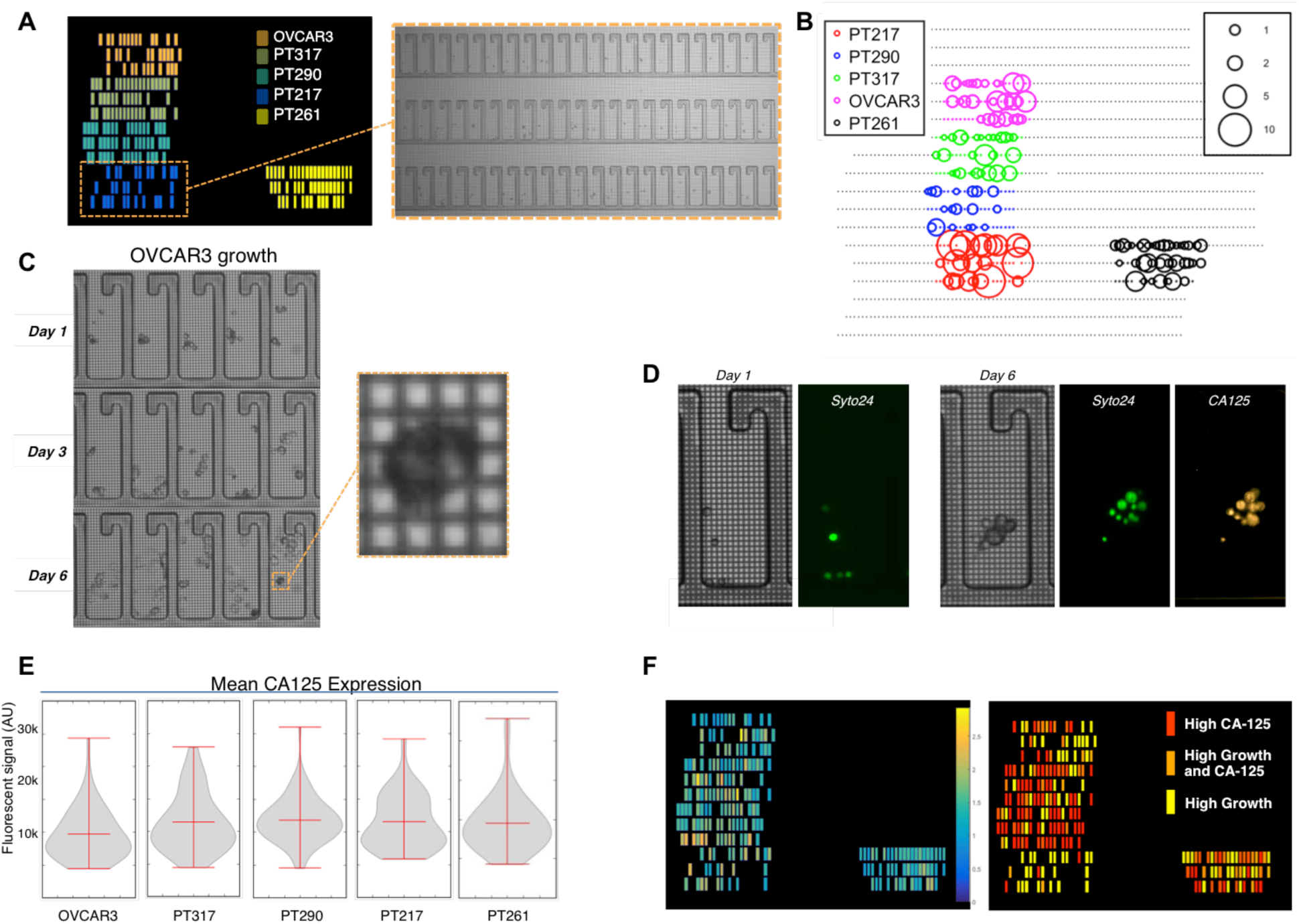
Phenotypic analysis of multiple ovarian cancer cell lines in a single chip. **(A)** Map showing penning location of 5 different ovarian cancer cell types, where each cell line is represented by a different color and brightfield image of cells in the selected region immediately after penning. **(B)** Cell growth of OVCAR3 and 4 patient-derived primary cancer cell lines overlaid on a map of the chip. Circle color indicates cell line, circle size correlates to cell growth factor in a pen (larger circles= larger growth factor after 6 days). **(C)** OVCAR3 ovarian cancer cells cultured on chip for 6 days, with spheroid formation. **(D)** Representative examples of brightfield and fluorescence images showing co-localization of Syto24 nuclei stain and CA-125 specific antibody. **(E)** Violin plots showing the distribution of single cell CA-125 expression for each cell line. **(F)** Heat maps showing average cellular CA-125 expression by pen (left panel) and average CA-125 multiplexed with cell growth by pen (right panel), demonstrating the ability of this approach to enable multiparameter phenotypic analysis.

Notably, given consistent culture conditions across the chip, each independent cell population has a unique set of growth kinetics. Tumor cells derived from PT217, who had stage IV cancer and succumbed to her disease 9 months after initial surgery, demonstrated the most aggressive growth rate, while those derived from PT290, who is currently the longest survivor in this cohort and presented with stage III disease, showed the slowest growth kinetics (patients were ranked by growth rate in decreasing order as follows: PT217, PT261, PT317 and PT290). By adjusting pen geometry, media conditioning, or starting cell number, the microenvironment can be optimized to provide robust growth conditions for the target cells. In particular, these data highlight several unique features of our approach which are the ability to simultaneously: 1) culture and characterize single or groups of patient-derived cells, 2) culture multiple cell types or mixtures, and 3) track and compare growth kinetics of individual cells within these populations.

### Single cell protein expression assays

The BLI platform also performs real-time monitoring of protein expression from independent cell populations over time with single cell resolution. In the context of ovarian cancer, the measurement of Cancer Antigen 125 (CA-125) levels is approved by the FDA as a proxy for monitoring ovarian cancer response to treatment and correlates with tumor burden.^26^ As such, it was of interest to investigate the ability to detect and measure CA-125 secretion at the single cell or subclone population level. Cell surface CA-125 expression was clearly detected and compared by fluorescent labeling from the four primary cell cultures and the immortalized cell line which had been multiplexed and cultured for 6 days on a single chip (Fig. 6D). A range of CA-125 expression levels was observed between patients and even among individual cells derived from a single patient, demonstrating the ability to monitor unique cell expression signatures over time. Co-localization analysis of multiple markers over time can be performed to assess biological function, as seen in Fig. 6D, which shows a high degree of overlap between Syto24 (a cell viability stain) and Anti-CA-125.

One significant advantage of this approach is that fluorescent signals are readily tracked and intensity values can be extracted on a per cell basis, organized by each unique pen. These differential expression results are encompassed in the violin plots in Fig. 6E, where the fluorescence of cells from the same population has been plotted, and each cell is represented individually. Statistical analysis of these data using the Kruskal-Wallis test shows that the CA-125 expression profiles are significantly distinct from one another (p < 0.0001). CA-125 fluorescence data was also plotted as a heat map to demonstrate the average CA-125 expression over all cells within one pen (Fig. 6F) and highlighted that expression levels varied by as much as 250 percent in both inter- and intra-cloned cell populations.

While the approach described here targets cellular CA-125 expression rather than CA-125 serum levels, it is interesting to highlight the concordance of these two metrics in PT217, who was a chemoresistant, stage IV HGSOC patient. This patient showed a unique CA-125 single-cell expression profile (Fig. 6E), which is distributed bimodally; these data suggest two distinct cellular subpopulations, including one with a high CA-125 expression level that is approximately double the other mean CA-125 values. Notably, PT217 also had, by over 3-fold, the highest CA-125 serum levels of all patients in this study (1578 U/mL versus 49 U/mL for PT261, 478 U/mL for PT290 and 187 U/mL for PT317).

In many cases, multiple phenotypic measures are needed to assess populations in cell culture, beyond just detection of a single protein marker. The heat map in Fig. 6F illustrates the integration of both growth and CA-125 expression. Integration of the two biologic characteristics demonstrates the appreciated fact that CA-125 expression levels do not correlate with growth rates across all cell lines, with the one exception that PT217 (whose disease was the most progressed and who had the highest CA-125 serum levels as discussed above) had a subpopulation of cells with the highest CA-125 expression levels as well as the highest growth rate.

### Single cell and bulk DNA sequencing from patient samples

As a demonstration of the downstream sequencing capabilities of this approach, we performed somatic variant screening on single cells derived from resected ovarian tumor tissue from a single patient, PT498, who had stage IIIC disease involving both ovaries and fallopian tubes. Furthermore, we compared the sequencing results to those from bulk tissue derived from the paired identical tumor using the BLI platform to isolate single cells as well as groups of cells for analysis. Single cell DNA was amplified using molecular displacement amplification (MDA), and the amplified material was sequenced using the Ion Torrent S5XL instrument to obtain high coverage data from the AmpliSeq HotSpot V2 panel, which was utilized for sensitive characterization of genomic variants across 50 oncogenic and tumor suppressor genes in each grouping of cells (single, 10, 100, 1000 or bulk, with the 10, 100 and 1000 groupings serving as controls). The batching function of the platform’s OEP technology facilitated the granularity to choose single cells or batches of cells for export off-chip for downstream molecular methods, sequencing, and analysis. This demonstrates the ability to select the appropriate sensitivity for the desired assay, as required by the heterogeneity of the biological question under investigation. To demonstrate this, 70 single- and 22 multi-cell samples were collected and comparative analysis of the detection of variants in those populations from the same tumor sample was performed. Fig. 7A is a dendrogram of the 73 cellular subsets passing filtering criteria ((n=51) single cell, (4) 10 cell, (8) 100 cell, (8) 1,000 cell and (2) bulk), showing independently sequenced single cell (or indicated cell grouping) variant profiles demonstrating the ability to identify genetic subclonality of this patient sample while conserving the remainder of the sample for orthogonal processes either on- or off-chip – filtering criteria are described in detail in Methods. Using these variant profiles, 5 major subclones/subclades with significant variance were identified across 9 oncogenes from the tumor of this patient as shown in Fig. 7 and Table 3. These data demonstrate the ability to use single cell genomics in tandem with the Beacon^TM^ prototype platform to detect low-frequency clonal variation that would otherwise be masked by major variants from the bulk tissue (i.e., yellow and green clade), recapitulating that no major DNA damage affecting targeted sequencing has occurred in cells selected using OEP. When the single cell variant profiles are compared to those derived from the 10-, 100- and 1000-cell sub-batches, the increased sensitivity provided by single cell sequencing is apparent.

**Figure 7.**
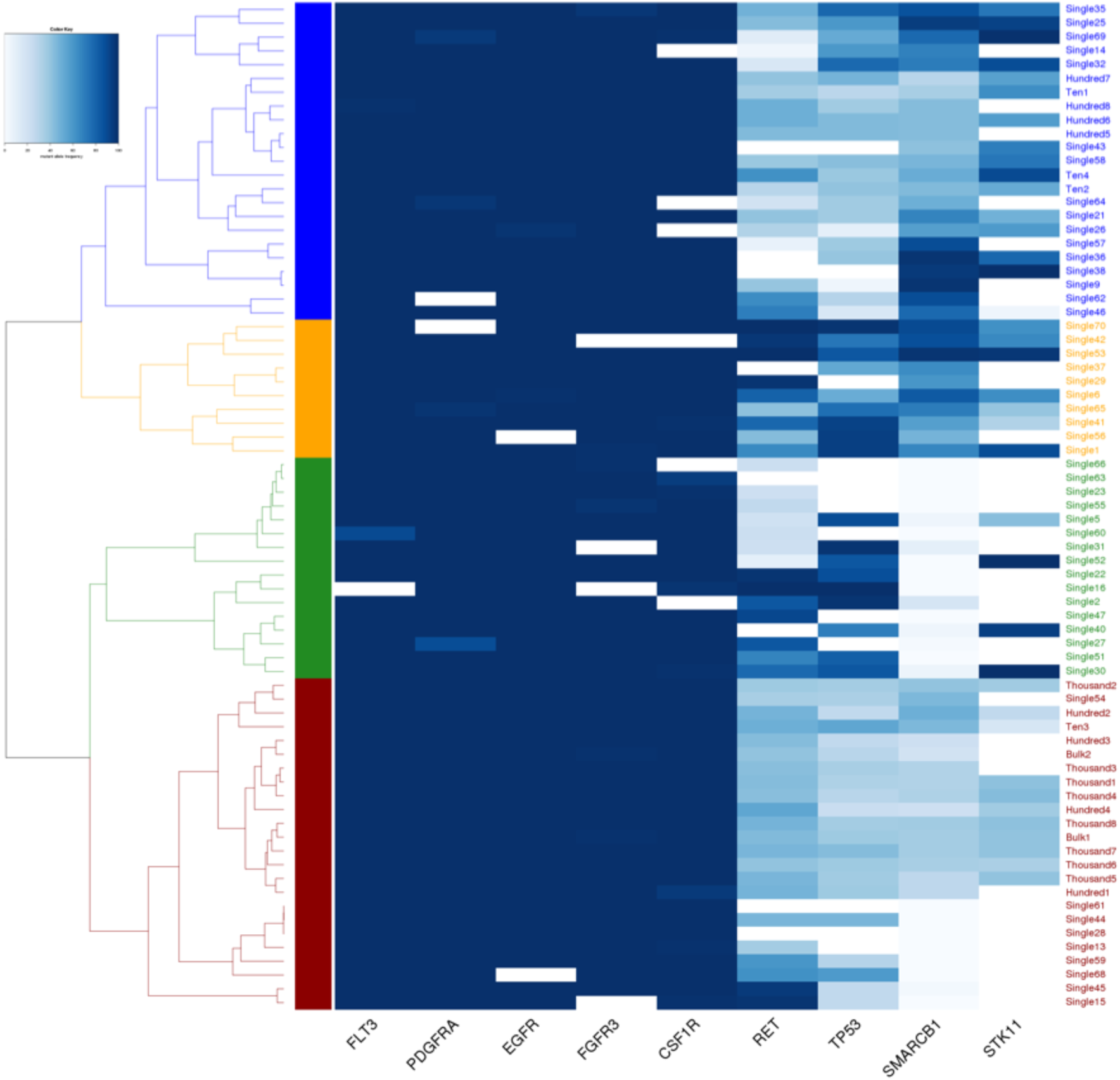
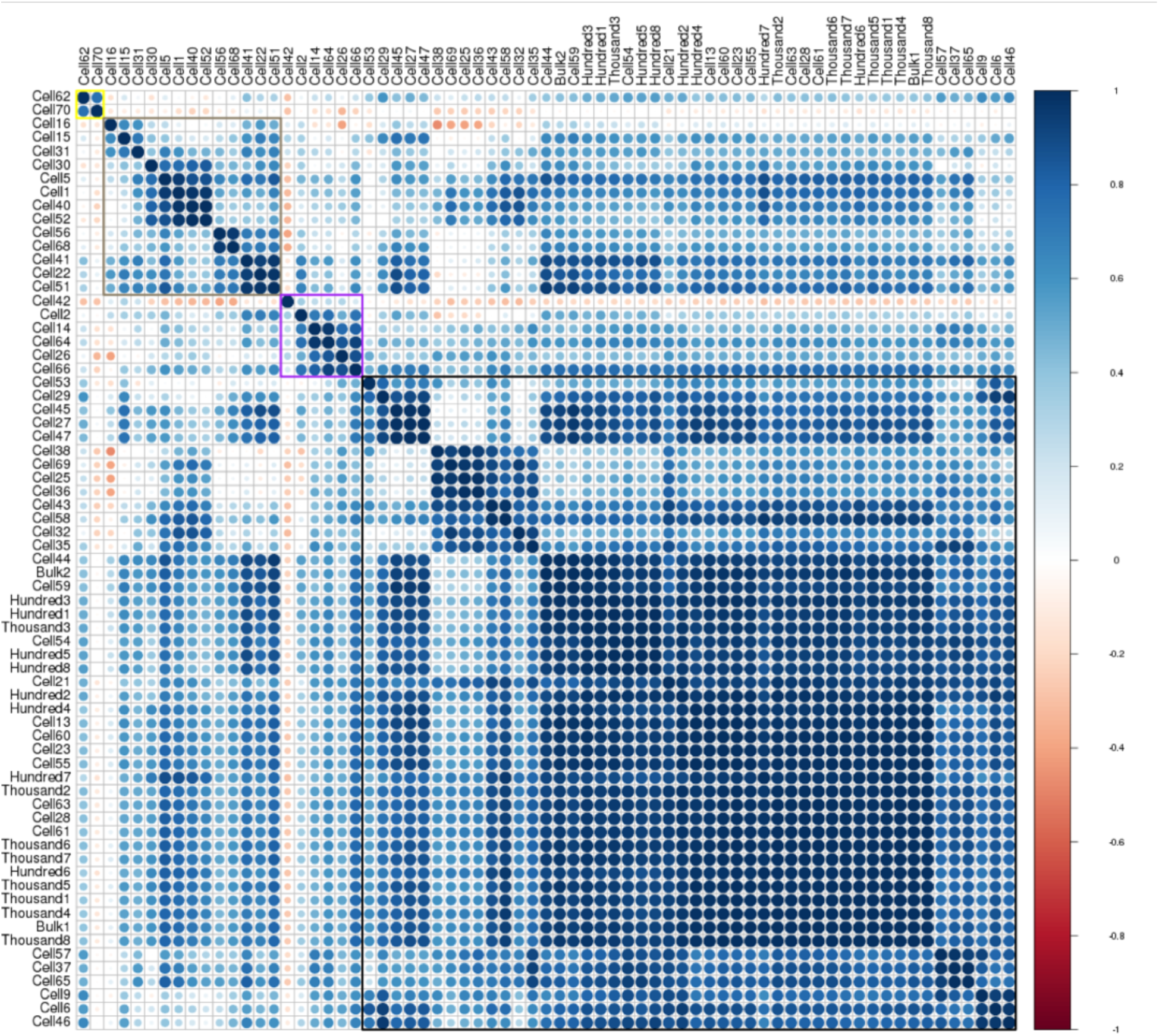
Allelle frequency estimates and cladding relationship from single and bulk cell genomic analysis. **(A)** Heat map showing color-coded mutant allele frequencies. Frequencies on a total of 9 variants depicted by the Genes (on X axis) they containing these mutations, were used from 73 samples, composed of 2 bulk samples, 8 hundred-cell samples, 51 single-cell samples, 4 ten-cell samples and 8 thousand-cell samples. The samples themselves (on Y-axis) are color-coded by the groups they define by the clades depicted by the dendrogram. **(B)** Correlation matrix ordered using the hclust method. The figure depicts positive correlations in blue and negative correlations in red. The color scale indicates estimated Pearson’s correlation coefficient values. Statistical significance in each correlation is proportional to the size of the circle. Groups of highly correlated samples, grouped by hclust method, are depicted using color-coded boxes.

Fig. 7B shows the single cell similarity plot comparing each individual cell or batched sample to reconstruct the subpopulations from the dendrogram as well as the concordance and discordance of all detected variants within the selected nine genes between all samples. This plot displays the associations between individual cells defining the subclades, but also highlights outlier cells (shown in red) within the subclade.

The ability to perform high-resolution DNA sequencing from these cells clearly demonstrates the applicability and robustness of this platform, which selected for single cells or cell pools and resulted in successful genetic variant profiling of a primary cancer tissue biopsy. The sequencing QC metrics validate the successful library construction and sequencing metrics from single cells based on mean read depth, on-target read percentage, and uniformity, relative to the bulk metrics, which are 1263X, 97.5% and 75%, respectively. The functional implications of the variants detected in these cell groups could be addressed in a continuation of this study using a larger oncogenic cohort and single cells isolated by the experimental platform for genomic profiling.

## DISCUSSION

In this publication, we demonstrate the multilayered and cross-functional phenotypic and genomic analysis of single cells made possible by the BLI Beacon^TM^ prototype platform. Using HGSOC as our primary model, we aimed to identify the potential of this platform for use in developing novel experimental pipelines to facilitate precision medicine in many fields, including oncology. This novel cell analysis system provides a powerful new approach to unraveling the signaling networks controlling the evolution of HGSOC disease, and potentially those of many other tumor types and disease states. Together, this study has demonstrated the ability of the experimental platform to select single cells with high accuracy and position them precisely for subsequent analysis. Applications include single- or multi-cell isolation, investigation of cell-cell interactions, microenvironmental studies for single cells, groups of cells or spheroids, quantitation of cell growth, single-cell temporal phenotypic analysis and export for genomic interrogation. As the system continues to be developed, the addition of additional analysis pipelines is anticipated.

Using HGSOC as a relevant pathologic system, single cell isolation also allowed for precise genomic analysis and a more accurate estimation of subpopulation frequency and status (e.g., homozygous normal, heterozygous, homozygous variant) in the cells and tissues examined here. Given that TP53 mutations are consistently present in ovarian tumor samples, it is not surprising that TP53 mutations are present in the two bulk samples from PT498.

Notable is that the allele frequencies measured in these bulk samples are 37.1% and 29.2%. Nonetheless, these allele frequencies are concordant with other studies utilizing standard sequencing approaches.^21^^,^^27^^,^^28^ It is of direct interest therefore to our approach of characterizing single cells and not just tumor in bulk. Examining 51 single cells yields more complex data; while 19 (37%) single cells had TP53 mutation frequencies consistent with bulk samples (25-75% frequency), 16 (31%) single cells had a much higher (>75%) TP53 mutation frequency and 16 (31%) single cells had a much lower (<25%) TP53 mutation frequency. These observations define the existence of subpopulations of TP53 mutated cells within the bulk tumor sample. Moreover, this approach enabled the detection of low frequency subclones that would otherwise be completely masked by major variants within bulk tissue variants (e.g., yellow and green clades in Fig. 7A). Strikingly, the TP53 mutation identified is the well-studied, relatively common SNP (rs1042522) resulting in either arginine or proline at position 72 (R72P). While a large number of epidemiologic studies have demonstrated mostly conflicting results on the association of this germline SNP with susceptibility for developing various cancers and survival, functional studies have identified numerous biochemical and biologic differences between the encoded proteins.^29^^,^^30^ To our knowledge, this is the first study identifying an active change in SNP frequency between tumor subclones within an individual, which strongly demonstrates the utility of the BLI platform and single cell analysis in general. The biologic relevancy of this change and its effect on subclonal populations of tumor cells will require future studies using an expanded sample cohort. An example of a low-frequency variant identified in this study, *SMARCB1 c.1119-41 G>A*, is also identified as a mutation in COSMIC (<u>cancer.sanger.ac.uk</u>), which demonstrates the ability to detect known mutations at lower frequencies using single cell sequencing approaches afforded by this technology; in this study, 26 of the 51 single cells expressed the *SMARCB1 c.1119-41G>A* variant. A comparison of the bulk population total variant profile to the 100-cell sub-batch profile, the 10-cell sub-batch profiles and the single cell profiles demonstrated the increased sensitivity provided by single cell sequencing. In future studies it is possible that this type of genetic information, combined with drug susceptibility data, may enable more precise genotype to phenotype correlations.^31^

This method also allowed for the establishment of precise and diverse cell phenotypic assays at very high resolution (i.e. single cell) and scale, which would increase the analytic power of experiments using limited or precious cell samples. It is especially notable that tumor cells derived from PT217 who was a chemoresistant, stage IV HGSOC patient had, by over 3-fold, the highest CA-125 serum levels of all patients in this study (1578 U/mL versus 49 U/mL for PT261, 478 U/mL for PT290 and 187 U/mL for PT317) and showed both a high CA-125 expression peak on the single-cell violin plots (Fig. 6E) and aggressive growth kinetics (Fig. 6B). It is clear from the CA-125 expression trends in Fig. 6E and 6F that only a subset of the cellular population exhibited this specific phenotype, which highlights the importance of a single-cell approach. The platform has the capacity to run thousands of precisely controlled experiments in parallel and measure the results rapidly, all on a single chip, providing a technical foundation upon which additional methods can be developed in order to analyze cellular heterogeneity. We envision that this type of approach could ultimately enable phenotypic screening of isolated tumor cells before the selection of the optimal patient therapy, replacing or enhancing other functional models.^32^^-^^34^ Testing a tumor at the single cell level would provide insight into whether a specific targeted agent will kill all cells in the sample, and just as importantly, rule out that a prospective treatment may allow a small subset of resistant cells to survive and develop into new tumors. We believe the only way to enable such a study is to use a platform with full workflow automation such that this type of screening and analyses could be performed on thousands of cells in parallel to ensure rare populations are not missed.

Increased scale and single cell resolution of tumor cell analyses has the potential to enable more accurate assessment of phenotypic and genetic heterogeneity in HGSOC.^33^^,^^35^ Greater accuracy will also enable a better understanding of the underlying biological drivers of the disease state and hopefully enable more precise diagnosis and treatment. This study focuses primarily on the technology and methods that demonstrate the utility of the BLI platform to interrogate intratumor heterogeneity, which supports characterization of linked single-cell phenotypic and genetic information across a culture system. Associating multifaceted phenotypic and genomic analyses will facilitate a more thorough analysis of tumor heterogeneity and could rapidly drive precision oncology development for other tumor types. Building on the conclusions from this work, future studies will focus on more detailed analyses, including cell replication with and without known chemotherapies or cellular therapies in HGSOC.

## METHODS

### Patient enrollment and sample collection

All blood and tumor samples were collected in accordance with the Institutional Review Board (IRB) of the Icahn School of Medicine at Mount Sinai at the time of the patient’s surgery (GCO# 10-1166). Patient-derived cell lines were also generated under an IRB-approved protocol. Written informed consent was obtained from each enrolled patient prior to their debulking surgery. Samples used in this study were collected from patients over a three-year period, from 2013 to 2016. Patient PT498 was selected for our proof of concept genomics study. Standard histology and molecular analyses were carried out to determine presence of cancer, histological subtype, stage, and grade. An IRB-approved protocol was followed to review the electronic medical record for clinical information regarding CA-125 levels in sera.

### Dissociation of samples and preparation for single-cell applications

Tissue containing macroscopic disease, as verified by a clinical pathologist, was obtained and placed in a 50 mL conical tube containing DMEM (Thermo Fisher Scientific, Waltham, MA #11965) supplemented with 10% FBS (ATCC, Mannassas, VA #30-2020) and 1% penicillin/streptomycin (Thermo Fisher Scientific #15140122). Representative pieces of tissue, with grossly non-tumor containing portions excluded, ranged in size from 0.5-3.0 cm and were separated using a tissue scalpel for mechanical and enzymatic digestion. Tissue samples were processed using the GentleMACS system (Milteny Biotec, Germany #130-096-427) using the “37C_h_TDK_1” protocol for soft human tumors as per manufacturer’s recommendation. Following GentleMACS disruption, the samples were passed through 40-µm cell strainers and subjected to 15 minutes of RBC lysis using BD Lysis Buffer (BD Biosciences, Franklin Lakes, NJ #555899) diluted to a 1X concentration in sterile deionized water. Next, cells were re-suspended in cell culture medium, stained using trypan blue for dead cell exclusion and counted using the Cellometer Mini Cell Viability Counter (Nexcelom, Lawrence, MA). The cell suspension was then diluted as needed to provide for single cell pipeline processing and downstream sequencing at >50×10^3^ cells in >20 µL and a target concentration of 20×10^6^ cells/mL. Remaining cells were plated on a bed of irradiated mouse feeder cells to start the generation of patient-derived cell lines and, if there were sufficient cells available, cryopreserved in FBS containing 15% DMSO and 5 uM ROCK inhibitor (SelleckChem, Houston, TX #S1049). BLI Priming Buffer was prepared by diluting 100X Nautilus suspension reagent (Berkeley Lights Inc., Emeryville CA #520-00002) in DMEM containing 10% FBS and 1% penicillin/streptomycin. Cells for BLI analysis were spun down and washed twice in BLI Priming Buffer and finally re-suspended in BLI Priming Buffer to a final volume of 50 µL and at a concentration of 20×10^6^ cell/mL, when possible.

### Culture of control and PDCL cell lines

Non-adherent OKT3 hybridoma cells (ATCC #CRL-8001) were grown in Iscove’s Modified Dulbecco’s Media (IMDM; ATCC #30-2005) supplemented with 20% fetal bovine serum (FBS) (ATCC, #30-2020) according to manufacturer recommended cell culture conditions. A2780, an ovarian cancer cell line, (Sigma Aldrich, St. Louis MO #93112519) were grown according manufacturer’s recommended cell culture conditions. Jurkat cells (E6.1, Sigma Aldrich #88042803) were grown in RPMI 1640 media, supplemented with 2 mM Glutamine and 10% FBS, according to manufacturer recommended cell culture conditions.

### Culture of human ovarian cancer cell lines

Ovarian cancer (OvCa) patient-derived cell lines (PDCLs) were developed using a ROCK inhibitor and fibroblast feeder cell system and cultured in Corning 75 cm^2^ tissue culture-treated flasks.^36^ OvCa PDCLs were cultured in medium containing, per 1.0 L: 665 mL DMEM (Corning Life Sciences, Teterboro, NJ #10-013-CV), 75 mL FBS (Corning Life Sciences #35-011-CV), 10 mL penicillin/streptomycin (Corning Life Sciences #30-002-CI), 250 mL Ham’s F-12 Nutrient Mix (Thermo Fisher Scientific #11765070), 5 mg insulin (Sigma-Aldrich #I9278), 50 µg hydrocortisone (Sigma-Aldrich #H0888), 10.0 µg EGF recombinant human protein (Life Technologies, #10605HNAE), 250 µg amphotericin B (Thermo Fisher Scientific #BP264550), 8.4 µg cholera toxin (Sigma-Aldrich #C8052), and 5.0 µmol of ROCK inhibitor (SelleckChem #S1049). NIH:OVCAR-3 cells (ATCC #HTB-161) were cultured in Corning 75 cm^2^ tissue culture-treated flasks in DMEM medium supplemented with 10% fetal bovine serum and 5% penicillin/streptomycin. The feeder cells used were Swiss 3T3-J2 mouse fibroblasts, irradiated at 30 Gy (3000 rad) using a cesium source irradiator prior to plating in co-culture at a ratio of 10:1. Cell lines grown using the feeder cell system were passaged once or twice in these conditions until a tumor majority was observed, at which time a portion were transferred from the feeder cell system and into feeder-cell conditioned medium. Feeder cell-conditioned medium was prepared in bulk when cell line development was considered complete, at which point cell lines were independent of the feeder system, free of stromal cells and determined to maintain known patient tumor mutations.

### Preparation of cell suspensions for penning

Conditioned medium was prepared as described.^36^^,^^37^ Irradiated feeder cells (1.0 × 10^7^ to 1.5 × 10^7^) were plated in 175-cm^2^ tissue culture flasks in 30 mL of F medium. The medium was collected 3 days later and centrifuged at 1000 × g for 5 min at 4°C. The resulting supernatant was passed through a 0.22-µm pore-size Millex-GP filter unit (EMD Millipore, Billerica, MA #SLGP033RS). Conditioned F medium was frozen and stored at -80°C. Three volumes of conditioned F medium were mixed with one volume of fresh F medium; this mixture was supplemented with 5 µmol/L ROCK inhibitor before use.

### Isolation of mononuclear cells from whole blood

Human peripheral blood mononuclear cells (PBMCs) were isolated from 10 mL of whole blood using Histopaque (Sigma-Aldrich #10771) according to the manufacturer’s protocol. Briefly, whole blood is layered on room temperature Histopaque in a 15-mL conical tube and spun at 400 *g* for 25 min without brake. Following the spin, the mononuclear cell layer is removed and washed 3 times with PBS.

### Culture and staining conditions for automated cell penning

A2780 ovarian cancer cells were incubated with RPMI-1640 media (Thermo Fisher Scientific #11-875-119) supplemented with 2 mM Glutamine (Thermo Fisher Scientific #35050061) and 10% FBS (Seridigm, Radnor, PA, #97068-101) containing Hoechst (BD Pharmingen, San Jose, CA #561908; 1:1000 Dilution of 1 mg/mL stock) and MitoView Green (Biotium, Fremont, CA, #70054; 1:1,000 Dilution of 200 µM stock) for 60 min at 37°C. A peripheral blood mononuclear cell (PBMC) sample was stained with MitoTracker and Anti-CD8, PE-(BD Biosciences #562282 1:1,000 Dilution) for 60 min at 37°C. Both cell types were washed 3 times with cell culture media and re-suspended in 1 mL at a concentration of 1 x 10^6^ cells/mL in RPMI-1640, supplemented with 2 mM Glutamine, and 10% FBS media, according to the BLI manufacturer’s protocol, and placed into a 96-well plate in wells A1 and A2. Cells were imported into the chip pens using single (A2780 cell line) or multiple (PBMC) rounds of OEP, and images were collected automatically in 4 fields with various exposure times (see Results).

### Immunofluorescent staining to detect protein expression

Immunofluorescent staining was performed on mononuclear cells derived from whole blood or OvCa cell lines dissociated by trypsinization (0.25% for 5 minutes at 37°C) into single cell suspensions. Antibodies for EpCAM (VU1D9 Mouse mAb Alexa Fluor^®^ 594 Conjugate #7319), CD45 (Abcam [F10-89-4] Alexa Fluor^®^ 488 #ab197730), CK7 (Abcam [EPR1619Y] - Cytoskeleton Marker Alexa Fluor^®^ 488 #ab185048), and CA-125 (Clone M11 Thermo Fisher Scientific #MA5-12425) were diluted 1:50 in cell culture medium and cells were resuspended at 2×10^6^ cells/mL in 300 µL total volume. After 1 h of staining at 4°C, cells were spun down and resuspended in 30 µL to a final concentration of 20×10^6^ cells/mL and loaded onto the chip.

### Machine vision algorithms for automated cell characterization

Images were acquired in the desired channels every 2 hours and a BLI-proprietary algorithm was used to automatically detect and count cells by pen. Moreover, this algorithm was also used to measure cell size, X,Ycoordinates and fluorescence intensity by pen.

### Characterization of cell growth and protein expression

For all cell culture, BLI chips were primed using fresh or conditioned media pre-buffered with 5% CO2 for 1 hour. Cells from a non-adherent hybridoma cell line (OKT3) were seeded at 2×10^6^/mL in 200 µL in a 96-well plate and OEP-loaded into pens as single cells. CO2-buffered media was perfused through the chip at 0.01 µL/sec. Images were taken to track growth at distinct time points.

For primary cell growth assessment and automated counting, cells were pre-stained with the Syto24 (Thermo Fisher Scientific #S7559), a live cell nuclear stain, prior to loading in order to enable accurate cell counting of adherent cells. For pre-staining, cells were incubated for 15 min at room temperature in conditioned media containing a 1:30,000 dilution of Syto24 (to minimize viability impact) and 1:100 priming additive (Berkeley Lights Inc. #520-00002). Cells were re-suspended at 5-10×10^6^ cells/mL in 30 µL of conditioned media. Cells were loaded onto the BLI chip at 25°C and bulk-penned using a waveform generator peak-to-peak (WFG PTP) voltage of 2.1 V at 5 µm/s, targeting 6-10 cells per pen. Each cell line was loaded into the pens of a specific region of the chip followed by image capture in brightfield (50 ms) and FITC channels (200 ms). Cross-contamination between cell lines was estimated to be non-existent by visual inspection. Cells were then cultured at 37°C in 5% CO2-buffered conditioned media containing 1:100 bovine fibronectin (Sigma-Aldrich #F1141), which was perfused at 0.01 µL/s. After 24 hours, the perfusion media was switched to 5% CO2-buffered conditioned media without fibronectin and the bulk media was replaced every 48 hrs. Cells were cultured on the chip for a total of 6 days with image acquisition every 2 hours (exposure time of 50 ms).

To assess expression of CA-125, surface expression levels were measured by importing a dilution of 1:5 CA-125 primary antibody (Clone M11, Thermo Fisher Scientific #MA5-12425) in conditioned cell culture media, followed by incubation of 60 minutes to allow diffusion of the antibody into the pens. The unbound primary was flushed out of the chip with 250 µL of media at 1 µL/sec. Next, goat anti-mouse secondary antibody conjugated with TxRed dye at 1:100 dilution in cell culture media was imported, followed by a 60 min incubation to allow diffusion of the secondary antibody into the pens. Finally, additional Syto24 dye, diluted 1:100, was loaded and incubated for 30 min to enable staining of cell nuclei. Brightfield and fluorescent images (FITC 200 ms; TxRED 7500 ms) were captured and analyzed to obtain protein expression levels for each pen and cell.

### Microscopy image processing and analysis

Aligned images stacks of Brightfield, FITC (Syto24) and Texas Red (CA-125) channels over all fields of view were generated. Within each pen, the ImageJ multipoint selection tool was used to select each cell identified by Syto24 staining in the FITC channel. The grey value for these selections was also measured on the associated Texas Red image in the stack, yielding CA-125 fluorescence values for each cell, organized by pen. This data was then plotted in Matlab to generate the heat maps and violin plots shown.

### Single cell Whole Genome Amplification (WGA)

#### Plate set-up and storage

Cell sorting was performed using the Single Cell Analysis and Genomic (SCAG) tool and samples were exported onto a 96-well plate and stored at -20°C for long term DNA processing. Cells were grouped on the plate according to cell number; i.e. 70 wells contained a single cell exported from the BLI tool into 2 µL of BLI Nautilus buffer suspended in 20 µL of mineral oil, 4 wells contained 10 cells each, 8 wells contained 100 cells each and 8 wells contained 1000 cells each. 2 wells of bulk tumor cells were also included, each of which contained approximately one million cells.

Upon thawing, the entire plate was simultaneously processed in order to avoid multiple freeze-thaw cycles and to prevent any DNA damage. Half of the bulk sample was processed for DNA extraction, using the PureLink Genomic DNA Mini Kit (Thermo Fisher Scientific #K182001) and subsequently purified with 1X Ampure XP beads to concentrate the sample. Whole genome amplification (WGA) was performed using molecular displacement amplification (MDA) on the remaining samples as described below.

#### Single Cell WGA

WGA was performed using the QIAgen REPLI-g Single Cell Kit (QIAgen, Germantown, MD #150345) MDA method, according to the manufacturer’s instructions. Briefly, Buffer DLB was reconstituted with 500 µl nuclease free H_2_O and mixed well; and Lysis buffer D2 was prepared for 12 reactions using 3 µL (1M) DTT and 33 µL Reconstituted Buffer DLB. Lysis buffer D2 preparation was scaled according to the number of reactions needed. To bring samples up to the appropriate volume, 2 µL of phospho-buffered saline (PBS) was added to each sample, for a total volume of 4 µL. Next, 3 µL of Lysis Buffer D2 was added to each sample and the contents were mixed by pipetting and briefly centrifuged. This lysis step was incubated on a thermocycler set at 65° for 10 minutes, and then 3 µL of stop solution was added to each sample, mixed, centrifuged briefly, and placed on ice. Immediately following this, 40 µL of WGA Master Mix, which included 9 µL nuclease-free water, 29 µL REPLI-g reaction buffer, and 2 µL REPLI-g sc DNA polymerase per reaction, was added to each sample and mixed by pipetting. The WGA reactions were incubated at 30°C for 8 hours. The REPLI-g DNA Polymerase was then heat inactivated at 65°C for 3 minutes.

#### WGA product purification

After WGA chemistry was completed and polymerase inactivated, a 1X AMPure bead-based purification was performed on the amplification products. Briefly, 50 µL of beads were added to each WGA reaction and incubated at room temperature while shaking on a vortex mixer at 2000 rpm for 10 min. The reactions were then placed on a magnetic separation device for 5 minutes or until solution was clear, at which point the supernatant was removed and discarded. While the tubes remained on the magnetic separation device, the beads were washed twice with 100 µl 80% ethanol. Without disturbing the beads, this wash solution was carefully removed and discarded. Following the wash the beads were briefly centrifuged to collect any residual ethanol from the tube walls, which was then removed and discarded. The reactions were removed from the magnet and the beads air-dried for 2 minutes. 55 µL of Elution Buffer (EB) was added to each reaction and mixed thoroughly to re-suspend the beads. The reactions were vortexed at 2000 rpm at room temperature for 2 min. Following mixing, each tube was spun down and placed on the magnetic separation device for 1 min or until the solution was completely clear. The eluate was removed and stored in a clean tube. The quantity and the quality of each WGA reaction product was determined using the DNA Broad Range Qubit Assay (Thermo Fisher Scientific #Q32853) and the 2100 Bioanalyzer, using the DNA 12000 kit (Agilent, Santa Clara, CA #5067-1508), respectively. Expected yield for each reaction was between 400-900 ng/µL. Samples were diluted to 10 ng/µL in EB and submitted for sequencing on the Ion Torrent platform. Multiple WGA technologies were assessed for false positive and negative rate, relative to the Qiagen Repli-g kit and deemed less efficient and more error prone, relative to Repli-g (data not shown).

### Ion Torrent library preparation and sequencing

#### Library Preparation

The Ion AmpliSeq Cancer Hotspot Panel v2 (Thermo Fisher Scientific #4475346) is a pool of multiplexed primers that amplify 208 “hot spot” areas of frequent mutation in 50 oncogenes and tumor suppressor genes. This panel was selected to amplify the DNA for library preps because it covers a broad range of genomic areas associated with cancer while allowing cost-effective multiplexing and is integrated into the Ion Torrent sequencing platform. Post-WGA, sample DNA was diluted to 10ng/µL and libraries were prepared with the Ion AmpliSeq Library Kit 2.0 (Thermo Fisher Scientific #4475345) using Ion Xpress Barcode Adapters 1-96 Kit (Thermo Fisher Scientific #4474517) to barcode each single cell, group of pooled cells or bulk tumor, following standard manufacturer’s instructions. Briefly, 30 ng of each sample were amplified using the Ion Torrent Cancer Hotspot panel and specified Ampliseq cycling conditions for the 208 amplicon pool. Following PCR amplification, the primers were partially digested with the proprietary FuPa enzyme and each sample was barcoded with a unique IonExpress barcode (1-96). Finally, a 1.5X bead purification was performed with Agencourt AMPure XP Reagent (Beckman Coulter, Brea, CA, #A63880) following the instructions in the Ion Ampliseq Library Prep protocol to clean up the sample and remove adapter dimers. All samples were quantified with the Ion Library TaqMan Quantitation Kit (Thermo Fisher Scientific #4468802) using a 1:100 dilution of each library in a 10 µL reaction volume, with 2 technical replicates per sample.

#### Priming the Ion Torrent sequencing chip

Following quantification, all samples were individually normalized to 100 pMol and 2 µl of each normalized library were combined to form a pool of 96 uniquely barcoded samples at a final concentration of 100 pMol. 25 µl of this pool was used for priming the sequencing chip. Ion Torrent chip priming and sequencing were carried out using the Ion Torrent S5XL system and the Ion Chef instrument with reagents from the Ion 540 Kit-Chef (Thermo Fisher Scientific #A27759). Briefly, the Chef was used to bind each library DNA fragment to Ion Sphere Particles (ISPs) and clonally amplify each fragment by emulsion PCR. Amplified DNA fragments were then bound to streptavidin-coated beads and template negative ISPs were washed away. Template-bound ISPs were then prepared for sequencing by loading onto one Ion Torrent S5 540 chip for sequencing on the Ion S5XL sequencing system.

#### Sequencing on the Ion Torrent Platform

The primed 540 chip was sequenced on the Ion Torrent S5^TM^ XL System with library read length set at 200 bp and 520 flows per chip, with all other instrument settings set to the manufacturer’s default for the Ion 540 Kit. Analyses of sequencing raw data were performed with Ion Torrent Suite (version 5.0.2) using the “coverageanalysis” and “variantcaller” plugins (with somatic/low stringency settings for the “variantcaller”), with all other settings for the run report set to the manufacturer’s default.

### Single cell genetics data analysis

#### Filtering of sequencing data

Raw reads were aligned to the hg19 reference human genome using tmap on Torrent Suite (TS). Only samples with ≥300X coverage, an on target value ≥90% and a uniformity ≥60% were further analyzed. Using these very stringent criteria, 51 of 70 single cell samples, 4 of 8 10-cell samples, 7 of 8 100-cell samples and 8 of 8 1000-cell samples were used for our analyses from PT498. In addition to the various cell pools, DNA extracted from bulk cells exported from the BLI instrument acted as a positive control.

#### Variant calling and filtering for bulk sample control analysis

Variants were called using the variant caller plug-in from the Torrent Suite^TM^ (Thermo Fisher Scientific) version 5.0.4 using default settings. In addition to standard homozygous and heterozygous variant calls, Torrent Suite also reported “Absent” and “No call” results for variants of interest from the targeted Hotspot panel. Variants present in ≥ 50% of each of the 10-, 100-, 1000-cell pools were used to create Virtual Bulk (VB) variants for each of the pools. VB variants of each pool were compared to the variants identified in the single cells. If a VB variant was also found in the single cell group at a frequency < 10%, this variant was filtered out, as it was more likely to have been introduced during WGA. In one instance, a FBXW7 variant (in red) was found in three pools was filtered out, as it was only observed at a frequency of <5% in the single cell samples. For the single cell group, variants present at ≥ 20/51 samples were categorized as VB variants.

#### Clustering of variants and cells with intra-single cell comparison

Variant frequencies for all samples, used in the analyses, for 9 genes were supplied to the heatmap.2 function in R (v 3.3.1) to generate the heatmap. Distance matrix was computed using dist() function in R (v 3.3.1). Variant frequencies for all samples, used in the analyses, for 9 genes were supplied to the corrplot() function in R (v 3.3.1). Variants and cells were distributed and clustered using a complete-linkage clustering algorithm, which was implemented as a standard function (hclust) in R.

### Data availability

All data that support the findings of this study are available from the corresponding author upon reasonable request.

## ACKNOWLEGEMENTS

We would like to acknowledge Magali Soumillon, Hayley Bennett, Thomas Vetterli and Debjit Ray for their technical assistance.

